# Detection of clustered anomalies in single-voxel morphometry as a rapid automated method for identifying intracranial aneurysms

**DOI:** 10.1101/2020.07.22.216812

**Authors:** Mark C Allenby, Ee Shern Liang, James Harvey, Maria A Woodruff, Marita Prior, Craig D Winter, David Alonso-Caneiro

**Affiliations:** Queensland University of Technology (QUT), Biofabrication and Tissue Morphology Group, Centre for Biomedical Technologies, Institute of Health and Biomedical Innovation, Kelvin Grove, Qld 4059, Australia; University of Queensland (UQ), Centre for Clinical Research, Faculty of Medicine, Herston, Qld 4006, Australia; Department of Medical Imaging, Royal Brisbane & Women’s Hospital, Herston, Qld 4029, Australia; Kenneth G Jamieson Department of Neurosurgery, Royal Brisbane & Women’s Hospital, Herston, Qld 4029, Australia; Queensland University of Technology (QUT), Contact Lens and Visual Optics Laboratory, Centre for Vision and Eye Research, School of Optometry and Vision Science, Kelvin Grove, Qld 4059, Australia

**Keywords:** Cerebral angiography, Intracranial aneurysm, Computational anatomy, Statistical shape analysis

## Abstract

Unruptured intracranial aneurysms (UIAs) are prevalent neurovascular anomalies which, in rare circumstances, rupture to create a catastrophic subarachnoid haemorrhage. Although surgical management can reduce rupture risk, the majority of IAs exist undiscovered until rupture. Current computer-aided UIA diagnoses sensitively detect and measure UIAs within cranial angiograms, but remain limited to low specificities whose output requires considerable neuroradiologist interpretation not amenable to broad screening efforts. To address these limitations, we propose an analysis which interprets single-voxel morphometry of segmented neurovasculature to identify UIAs. Once neurovascular anatomy of a specified resolution is segmented, interrelationships between voxel-specific morphometries are estimated and spatially-clustered outliers are identified as UIA candidates. Our automated solution detects UIAs within magnetic resonance angiograms (MRA) at unmatched 86% specificity and 81% sensitivity using 3 minutes on a conventional laptop. Our approach does not rely on interpatient comparisons or training datasets which could be difficult to amass and process for rare incidentally discovered UIAs within large MRA files, and in doing so, is versatile to user-defined segmentation quality, to detection sensitivity, and across a range of imaging resolutions and modalities. We propose this method as a unique tool to aid UIA screening, characterisation of abnormal vasculature in at-risk patients, morphometry-based rupture risk prediction, and identification of other vascular abnormalities.

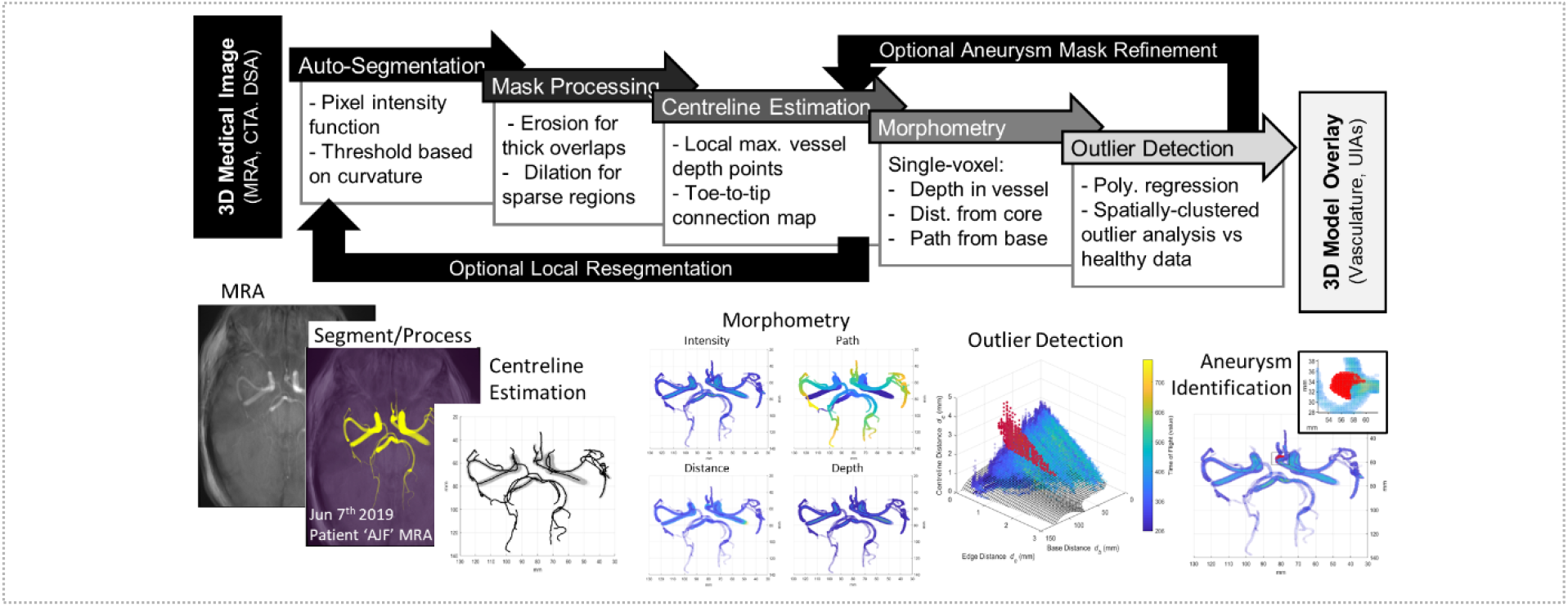
Graphical Abstract.

**Highlights:**

- Rapid and automated detection of unruptured intracranial aneurysms (UIAs) in MRAs
- Highly specific, sensitive UIA detection to reduce radiologist input for screening
- Detection is versatile to image resolution, modality and has tuneable mm sensitivity

## 1. Introduction

Intracranial aneurysms (IAs) are bulging, weak outpouchings of arteries that supply blood in the brain. IAs are relatively common (2% to 5% prevalence; Figure 1A,B) but rarely discovered prior to incident (International Study of Unruptured Intracranial Aneurysms (ISUIA), 2003). While most unruptured IAs (UIAs) are asymptomatic, between 0.25% and 1% spontaneously rupture resulting in a subarachnoid haemorrhage, a catastrophic event associated with a 40 to 50% mortality rate and with 50% of survivors left with permanent disabilities (Figure 1C) (Leng et al., 2018; Thompson et al., 2015; van Gijn et al., 2007; Williams and Brown, 2013). Therefore, early detection of UIAs is paramount so that management to prevent future rupture can be considered (Figure 1D,E) (Mayo Foundation, 2017).

**Figure 1:**
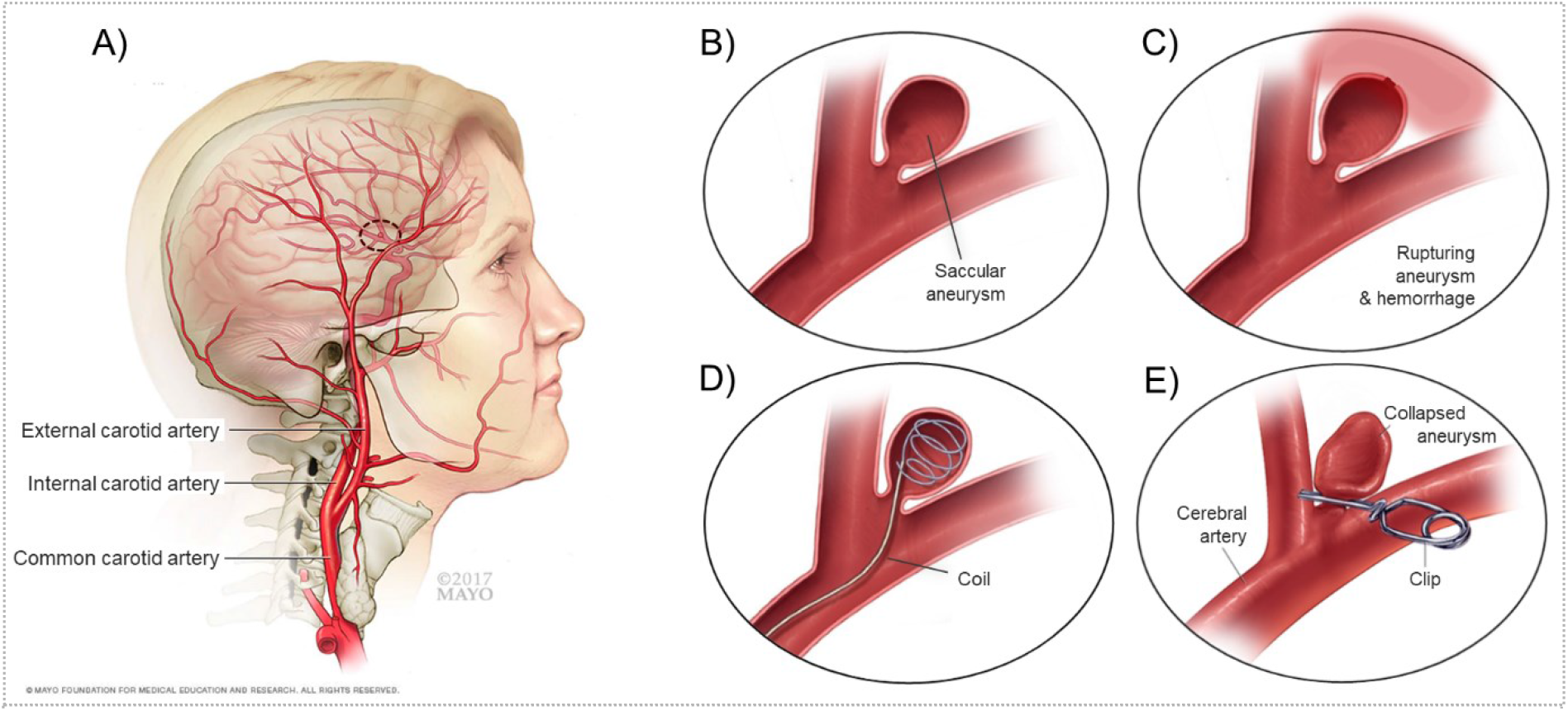
Intracranial aneurysm pathology and treatment. (A,B) IAs are bulging vessels within the head that are at risk of (C) rupture, causing subarachnoid haemorrhaging. Incidentally discovered UIAs can be monitored or surgically treated through (D) endovascular coiling or (E) neurosurgical clipping, which both carry nontrivial risk. *Adapted from the Mayo Foundation*.

Aneurysms are detected and measured by radiologists interpreting computed tomography angiograms (CTA), digital subtraction angiograms (DSA), or magnetic resonance angiograms (MRA) (Li et al., 2009; Okahara et al., 2002). In 65% to 91% of cases UIA detection is incidental, and the frequency of incidental detection has been rising due to an increased use of high-resolution intracranial MRAs which do not require intravenous contrast or x-ray radiation (Corfield et al., 2010; Duan et al., 2018; Mair, 2015; Thompson et al., 2015). However, incidental detection is often by radiologists not specialised in neuroanatomy who are not searching for an UIA. Therefore, detection sensitivity may be limited by the shortage of experienced neuroradiologists able to review the increasing number of cranial radiology examinations (Z. Shi et al., 2020).

Computational analyses can provide a rapid and automated identification of UIAs to serve as a supportive reference to the nonspecialist radiologist for increased UIA detection accuracy. As radiologist UIA detection sensitivity can be as low as 64% (Miki et al., 2016), computer-aided diagnoses are attractive to improve accuracy while increasing the number of patient images analysed. Conventional computer aided diagnoses detect UIAs using predetermined morphometries of interest (curvature, sphericity, convexity), but have been less able to accurately identify irregular or small UIAs (Jin et al., 2016; Yang et al., 2011). Recent machine learning approaches surmount conventional limitations by detecting nonintuitive patterns consistent in aneurysmal regions, however their performance must be conditioned on large annotated training sets often unavailable for rare incidentally discovered UIAs. To date, conventional and machine learning approaches have achieved above 80% sensitivity but at the cost of low specificity, generating 4 to 41 additional false positive UIAs per image (Faron et al., 2019; Jin et al., 2016; Nakao et al., 2018; Stember et al., 2019; Ueda et al., 2019), which may not reduce the time required during radiologist interpretation (Z. Shi et al., 2020).

Herein, we present an image analysis method based on a single-voxel morphometry approach to rapidly, automatically, and specifically identify UIAs. Our algorithm automatically reconstructs 3D presentations of patient neurovasculature from MRA, CTA, or DSA image datasets, then seeks to identify UIA candidates based on three voxel morphometry attributes: distance from vessel centreline, distance from vessel edge, and distance from image base. Identified UIA location and size are measured and validated against clinician measurement. We analysed a cohort of 29 TOF MRAs presenting UIAs who are benchmarked against 705 healthy TOF MRAs. Our automated algorithm is unique in its rapid analysis of large 3D datasets without the need for training data or interpatient comparisons, its high and tuneable specificity and sensitivity to identify fine features for clinical observation, and its versatility in analysing a range of MRA resolutions as well as CTA and DSA modalities.

## 2. Methods and Calculation

Medical imaging records of 14 patients exhibiting at least one unruptured intracranial aneurysm and 27 healthy patients were retrospectively recruited from 2009 to 2019 and de-identified by the Medical Imaging and Neurosurgery Departments of the Royal Brisbane & Women’s Hospital according to ethical clearances (LNR/2019/QRBW/49363). While each healthy patient was imaged once, many patients harbouring UIAs were monitored over several years so that a total of 29 aneurysm-containing TOF MRAs were imaged. The UIA parent vessel, location, and dimensions were described within radiology reports and annotated within 2D slices of the image as illustrated in Figure S1. A further 678 healthy TOF MRAs were acquired from publicly available repositories (MIDAS and IXI) as described previously (Mouches and Forkert, 2019). TOF MRA, CTA, and DSA images were captured in the transverse direction at slice XY-resolutions spanning [0.18 × 0.18] to [0.61 × 0.61] mm/pixel and slice steps with Z-resolutions from 0.38 to 2 mm/pixel.

A Dell Latitude 5300 laptop with 16 GB RAM and 1.90 GHz Intel Core i7-8665U processor produced all timed runs. The proposed MATLAB algorithm (The MathWorks Inc, Natick, USA) resembles a pipeline with several distinct steps, as illustrated in Figure 2 and detailed in the following subsections.

**Figure 2:**
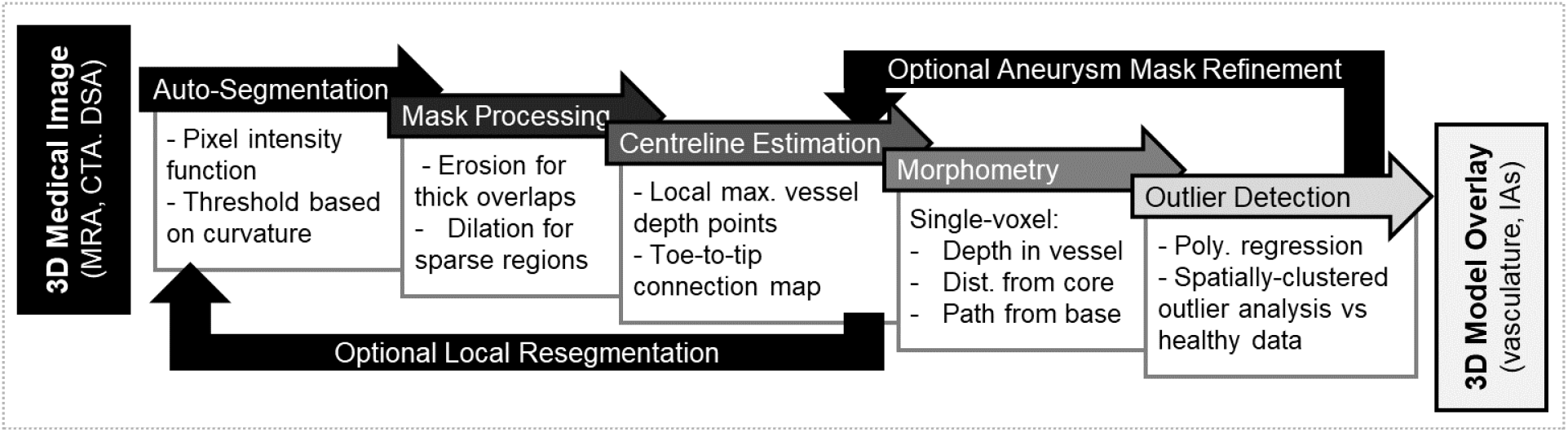
Intracranial aneurysm identification algorithm pipeline. The algorithm can input several different image modalities, including MRA, CTA, DSA. The images first undergo a global **Auto-Segmentation** to generate 2-3 tissue masks, the segmented neurovascular mask undergoes several Mask Processing steps, followed by a **Centreline Estimation** throughout the mask. Using the centreline, several **Morphometry** metrics are applied to measure geometrical properties of each voxel within the segmented vascular mask. The final **Outlier Detection** step identifies voxel properties consistent with normal vasculature, and regions which have large numbers of abnormal voxel properties. These regions are segmented as aneurysmal candidates for clinician assessment and computational measurement.

### 2.1. Global segmentation of a neurovascular mask (*Auto-Segmentation*)

Initially a universal threshold value was calculated to segment intravascular blood from surrounding cranial tissue (Figure 3A). All voxel intensities throughout the CTA, DSA, or TOF-MRA image were linearly divided into *n*_*bin*_ = 50 bins from maximal to minimal voxel intensity, similar to Nyúl et al., 2000. Voxel intensity brightness, *x*, belonging to different tissue types, such as dim extravascular soft tissue *versus* bone *versus* bright intravascular blood, was estimated by a sum of lognormal distributions (Figure 3B) as previously performed for MRA images (Forkert et al., 2012):

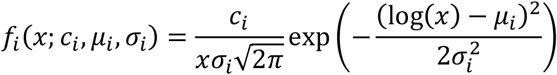

With tissue content scalar *c*_*i*_ and tissue intensity mean *μ*_*i*_ and standard deviation *σ*_*i*_. Voxel intensities throughout the image were estimated as the sum of these lognormal distributions. In CTA images all 3 aforementioned tissue types could be detected simultaneously, *f*_1+2+3_, while in TOF MRA and DSA images only intravascular and extravascular tissue could be distinguished, *f*_1+2_. The six or nine parameters of *f*_1+2_ or *f*_1+2+3_ were fit by minimising normalised error to the log voxel intensity of histogram bin 3 to 48 as to avoid overexposed and underexposed inconsistencies (removing *E*_*bin*_ = 4% of bins from either end). A threshold value, *T*_*v*_, was calculated to segment vessels from the brain tissue:

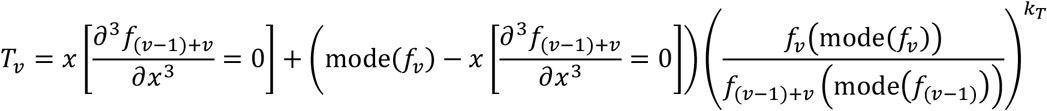

Where *v* is the vessel mask index (*v* = 2 for TOF MRAs and DSAs, *v* = 3 for CTAs), where 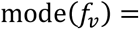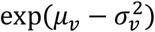 represents the mode of lognormal distribution *f*_*v*_, and where 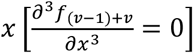 describes a critical point of inflection change between the modes of *f*_*v*_ and *f*_*v*−1_ as shown in Figure 3B. Explicitly determined weight parameter *k*_*T*_ dictates neurovascular mask size, later leveraged to normalise segmentation across different MRA resolutions.

**Figure 3:**
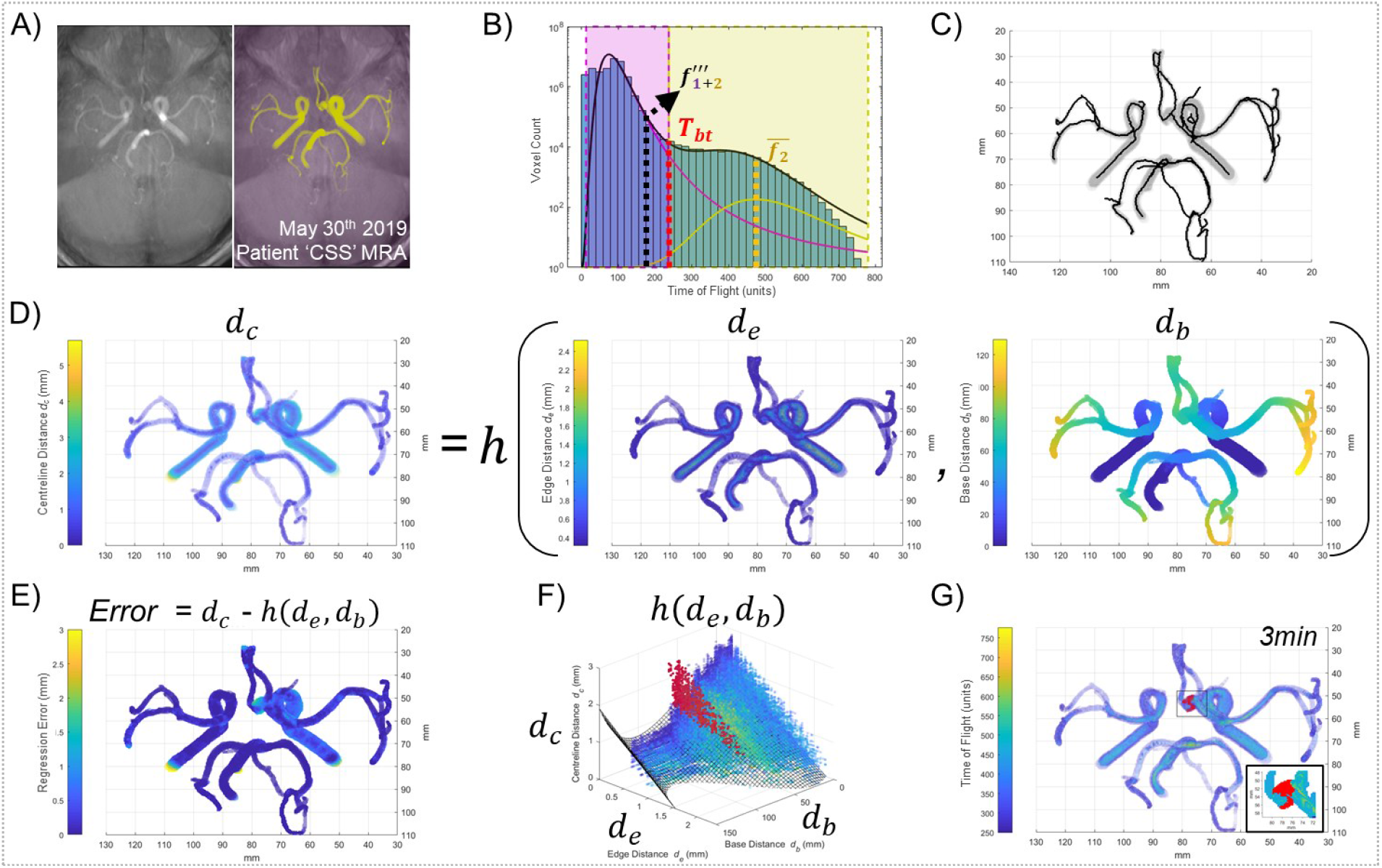
Intracranial segmentation, vascular network analysis, and aneurysm identification. (A,B) Segmentation of a 3D neurovascular mask based on a sum of lognormal distributions. (C) Identification of centrepoints via mask erosion, connecting adjacent centrepoints bottom-to-top into a network of branches, and measuring branch lengths and bifurcation points to detect centrepath length. (D) A voxel-specific polynomial regression was individually fit to measure the correlation between voxel distance from centreline (radius) versus voxel distance from vessel edge (depth) and voxel distance from base (path) for each image. (E) The error of actual voxel radius from expected voxel radius was measured and (F) outliers which were spatially-clustered together were identified as aneurysmal candidates, (G) overlaid in red over segmented mask image intensities with UIA region inset. The algorithm required 3 minutes to execute.

### 2.2. Evaluating single-voxel morphology (*Mask Processing* and *Centreline Estimation*)

Once the neurovascular mask is segmented, centrepoints are defined *via* a 3D medial surface thinning operation after filling closed voxel holes in the image. These centrepoints were then dilated and annealed together using a *m*_*dil*_ = 2 voxel rolling ball, and then centrepoints were re-identified in order to remove overestimated and overlapping small vessels using a range of binary morphological operations. A centreline was then connected through neighbouring centrepoints starting with the centrepoint in the lowest (most caudal) axial plane. When the next nearest unallocated centrepoint was further than *d*_*link*_ = 1mm away, the terminal end of that vessel branch had been reached and the centreline for a new vessel was evaluated from the next lowest centrepoint. Branch endpoints between *d*_*link*_ = 1mm and *d*_*latch*_ = 2mm away from a previously allocated centrepoint were then connected to form the full neurovascular tree. Once completed, an interconnected branch network of centrelines through the segmented neurovascular mask was formed (Figure 3C).

Voxels within the vascular mask were evaluated using a range of morphometric properties: taking the vascular voxel as reference, *d*_*c*_ indicates the distance to vessel centreline, *d*_*e*_ distance to edge, and *d*_*b*_ distance to the carotid vessel, defined as the nearest terminal centrepoint at the bottom of the mask (Figure 3D). The first two properties were calculated using Euclidean distance operations performed on Boolean matricies between the neurovascular model and points defining the vessel centrelines. The third distance operation required sorting individual vessel branches’ distance to the bottom of the vascular tree. Since many vessel branches intersect with many other vessel branches at multiple points, a combinatorial approach identified the tree branch path network which minimised total mask *d*_*b*_ across all intersecting branch entrances and exits.

### 2.3. Identifying aneurysms as outlier voxel clusters (*Morphometry* and *Outlier Detection*)

Each voxel’s three morphometries formed a consistent relationship. That is, the distance of a voxel from the vessel centreline (*d*_*c*_) depended on how far it was from the vessel edge (*d*_*e*_) as well as its distance away from the base of the mask (*d*_*b*_) (Figure 3E). A voxel far from the vessel centreline was more likely to be near the vessel edge and these distances were larger nearer the thick carotid arteries and not thin terminal cranial vessels. This relationship was characterised through fitting a polynomial regression through all voxels within the neurovascular masks (>10,000 voxels). For speed, a polynomial regression was used to estimate a voxel’s centreline distance (*d*_*c*_) which was first-order in its edge distance (*d*_*e*_) and fourth-order in its base distance (*d*_*b*_). Several other global and local polynomial regressions (loess) were compared but either exhibited a worse regression fit or prohibitively long processing times, respectively. Clusters of voxels which had centreline distances inaccurately predicted by our ℎ(*d*_*e*_, *d*_*b*_) regression were treated as outliers and considered to belong to either noncylindrical or inadequately large vessel anatomies, suggestive of vascular regions that may be aneurysm candidates (Figure 3F,E). Since this outlier detection is based on a single patient image and not a large dataset of patients, it can be particularly universal to anatomical and imaging differences.

The definition of these voxel cluster outliers, or aneurysm candidates, can be tailored for high or low detection sensitivity based on user demand and clinical application. For our multi-repository validation we identified aneurysm candidates as voxel clusters beyond the polynomial regression’s *E*_*out*_ = 96% confidence interval and larger than *V*_*min*_ = 17.3*h*(*d*_*e*_, *d*_*b*_) + 1.5 mm^3^ to allow for aneurysm detection in large central as well as small peripheral vessels. Voxel centreline distance estimated from *h*(*d*_*e*_, *d*_*b*_) typically varied from 1.5 mm at the carotid artery base to 0 mm at the terminal ends of peripheral arteries. We later demonstrate how users can decrease *E*_*out*_ and *V*_*min*_ to increase detection sensitivity for small or emergent aneurysmal buds.

### 2.4. Refining poorly-segmented neurovascular masks (*Optional Local Resegmentation*)

For poor quality images with under- or over-saturated cranial images, it may be difficult to analyse morphometry due to a fusing of adjacent vessels or an inaccurate capture of the aneurysm shape (Figure 4A). In such cases, a more accurate segmentation can be achieved through re-estimating the segmentation threshold for many small regions throughout the vessel structure. However, this more accurate local segmentation comes at the price of increased time and experimentally evaluated parameters. This local segmentation procedure proceeds by evaluating the vessel centrepoints produced from Section 2.2 and drawing spherical neighbourhoods around these centrepoints for re-segmentation. Next, successive centrepoints neighbourhood are considered until all their contained vessel masks have been re-segmented. Finally, the remaining image space is also re-segmented based on the threshold assigned to each voxel’s closest neighbourhood, which allows for significant mask growth.

**Figure 4:**
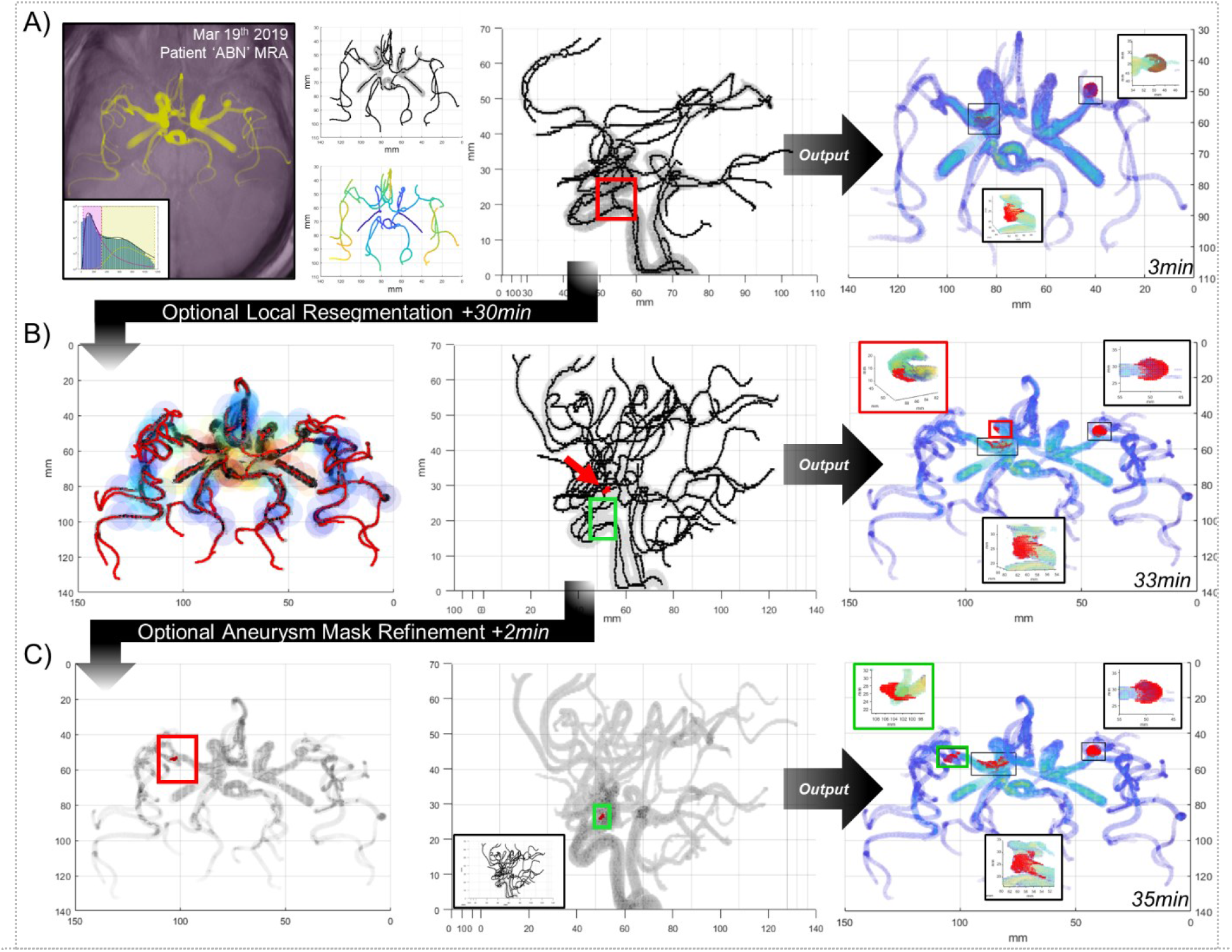
Optional refinements of the UIA identification pipeline. (A) While the algorithm pipeline presented in Figures 2 and 3 is rapid, sensitive, and specific, it struggles with large convoluted neurovascular masks with overlapping vasculature. When a more detailed mask and IA identification is required at the expense of increased computational time, additional pipeline refinements can be performed. (B) A local resegmentation can be performed which recalculates the threshold of the neurovascular mask within spherical neighborhoods along the previous centrelines, and then expands those thresholds into the remainder of the image. This resegmentation serves to cull vessel thicknesses within overlapping high-signal regions and enhance the vessel mask into regions with low signal and can be iterated as desired. (C) A mask refinement can be performed which surveys an aneurysmal candidate and deletes vessel centrelines which enter an aneurysm to provide a more complete IA segmentation.

The diameter of these spherical neighbourhoods must be large enough to consider the thickest vessel diameter while sampling frequency along the centrepoints must be fine enough to not allow unconsidered gaps. To ensure adequate neighbourhood overlap, sampling frequency was dictated by spherical neighbourhood centroid spacing scaled to sphere and vessel radius:

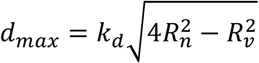

Where *R*_*n*_ is the desired spherical neighbourhood radius, *R*_*v*_ is the calculated vessel mask radius, and *k*_*d*_ is a user-defined scalar between 0 and 1 adjusting neighbourhood overlap, where *k*_*d*_ = 1 would allow neighbourhoods to overlap just enough to encompass the current vessel radius. Spherical neighbourhoods which are too large or too infrequent would limit the effectiveness of the local segmentation, but small or frequent neighbourhoods create unnecessary computational burden. Our masks appeared to work best with *R*_*n*_ = 2.75 mm and *k*_*d*_ = 0.5 to allow for re-segmented masks to grow in dim peripheral regions but also shrink in bright carotid regions (Figure 4B).

Spherical neighbourhoods can predominately be comprised of brain tissue near shrinkingly small vessels, resulting in only one lognormal distribution of low voxel intensities and skewing the calculation of a threshold value to extremely low values for re-segmentation, causing brain tissue to be interpreted as vessels. To avoid this, we ensure these neighbourhood minimum or maximum thresholds do not exceed predefined global limits. Specifically, we propose: *if T*_*v*_ < *T*_*min*_, *then* 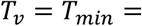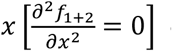 or *if T*_*v*_ > *T*_*max*_, *then T*_*v*_ = *T*_*max*_ = mode(*f*_2_), which represent the minimum and maximum values *T*_*v*_ is defined to have if two or more tissues are present.

### 2.5. Refining poorly-segmented aneurysmal masks (*Optional Aneurysm Mask Refinement*)

For several neurovascular masks harbouring an intracranial aneurysm, a network of vessel centrepoints determined by the binary erosion technique outlined in Section 2.2 could erroneously include centrepoints of the UIA leading to misidentification or an incomplete aneurysmal mask. To address this issue, we added a function which first identifies UIA candidates with increased sensitivity (*E*_*out,re*_ = 60%) as in Section 2.3, then for each UIA candidate determines whether a nearby branch of centrepoints terminates. Branches closer than *R*_*cull*_ from the UIA centroid and within *d*_*cull*_ from terminal centrepoint are culled. This process can be applied to trim many small branches diverging from major vessels by lowering other IA detection parameters such as *V*_*min*_ from Section 2.3. After the UIA-entering branches are deleted, the UIA detection is re-run at the original threshold sensitivity (Figure 4C).

## 3. Results

Our image analysis approach is conventional in specifying which morphometry attributes are indicative of UIAs but also mimics machine learning approaches by identifying unique patterns in single-voxel morphometries for each image. In doing so, our algorithm becomes adaptable to image quality and detection accuracy inputs. Our image processing steps require user-selected parameters (Table 1) which must be validated to be widely applicable but whose detection sensitivity can also be adjusted to detect fine or coarse aneurysmal candidate regions. Correspondingly, we validate the accuracy and demonstrate the versatility of our statistical approach herein.

**Table 1:**
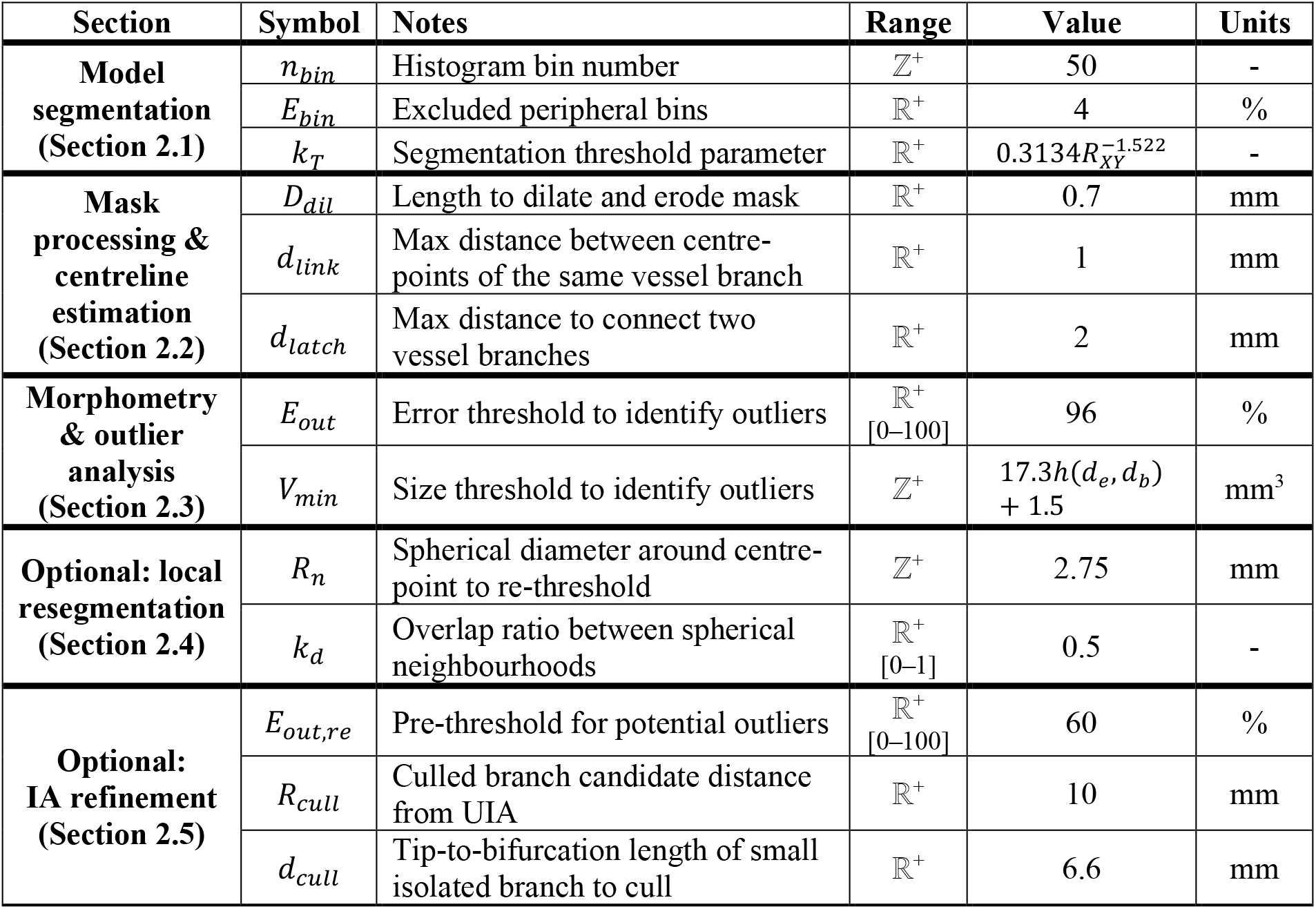
Summary of algorithm parameters. These algorithm parameters are grouped into the pipeline steps specified in Figure 2 and throughout methodology Section 2. Values are provided as a guide to replicate the algorithm performance in this paper but can be changed based on user preferences. Detection sensitivity was lowered in Figure 7 to demonstrate the impact of varied detection sensitivity.

| Section | Symbol | Notes | Range | Value | Units |
| --- | --- | --- | --- | --- | --- |
| **Model segmentation (Section 2.1)** | $n_{bin}$ | Histogram bin number | $\mathbb{Z}^+$ | 50 | - |
| | $E_{bin}$ | Excluded peripheral bins | $\mathbb{R}^+$ | 4 | % |
| | $k_T$ | Segmentation threshold parameter | $\mathbb{R}^+$ | $0.3134R_{XY}^{-1.522}$ | - |
| **Mask processing & centreline estimation (Section 2.2)** | $D_{dil}$ | Length to dilate and erode mask | $\mathbb{R}^+$ | 0.7 | mm |
| | $d_{link}$ | Max distance between centre-points of the same vessel branch | $\mathbb{R}^+$ | 1 | mm |
| | $d_{latch}$ | Max distance to connect two vessel branches | $\mathbb{R}^+$ | 2 | mm |
| **Morphometry & outlier analysis (Section 2.3)** | $E_{out}$ | Error threshold to identify outliers | $\mathbb{R}^+$<br>[0–100] | 96 | % |
| | $V_{min}$ | Size threshold to identify outliers | $\mathbb{Z}^+$ | $17.3h(d_e, d_b) + 1.5$ | mm<sup>3</sup> |
| **Optional: local resegmentation (Section 2.4)** | $R_n$ | Spherical diameter around centre-point to re-threshold | $\mathbb{Z}^+$ | 2.75 | mm |
| | $k_d$ | Overlap ratio between spherical neighbourhoods | $\mathbb{R}^+$<br>[0–1] | 0.5 | - |
| **Optional: IA refinement (Section 2.5)** | $E_{out, re}$ | Pre-threshold for potential outliers | $\mathbb{R}^+$<br>[0–100] | 60 | % |
| | $R_{cull}$ | Culled branch candidate distance from UIA | $\mathbb{R}^+$ | 10 | mm |
| | $d_{cull}$ | Tip-to-bifurcation length of small isolated branch to cull | $\mathbb{R}^+$ | 6.6 | mm |

### 3.1. Rapid automated IA detection is sensitive and specific

The sensitivity of this algorithm was validated over a cohort of patients presenting to the Royal Brisbane and Women’s Hospital (RBWH) between 2009 and 2019 who harboured an UIA imaged by TOF MRA (14 patients, 29 TOF MRA images). Several patients were imaged on more than one occasion to monitor aneurysmal shape changes over time and assess rupture risk for surgical decision-making. Accuracy of algorithm-identified aneurysm location and size was validated by interventional radiologists asked to retrospectively annotate, measure, and comment on unedited TOF MRA image slices while blinded to the algorithm’s detection as illustrated in Figure S1. The specificity of this algorithm was validated over a cohort of 27 patients not identified to have an intracranial aneurysm using TOF MRA and DSA. Additionally, the specificity of this algorithm was further validated over public IXI and MIDAS repositories for another 678 healthy TOF MRA patients (Mouches and Forkert, 2019). Images were automatically excluded from consideration if severe TOF MRA imaging artefacts interrupted the neurovascular mask construction (N = 6), or if only 1 lognormal distribution could be detected during segmentation (N = 94 to N = 154 depending on normalisation) so that a neurovascular mask could not be identified. This exclusion criteria and rate is consistent with prior publications (14% to 23% excluded *versus* 20% previously; Mouches and Forkert, 2019).

To provide a fair comparison, all validations were performed with identical parameters, which also ensured the algorithm was fully automated. The speed of the algorithm varied between 1.5 and 12 min primarily depending on image resolution (which affected image matrix size), and neurovascular mask volume (which affected time-intensive binary distance operations). The 734 TOF MRAs varied across 10 image resolutions from 0.26 to 0.80 mm/voxel (Figure S2). Median processing times per TOF MRA image were 5.2 min, 2.7 min, and 2.0 min for RBWH, MIDAS, and IXI datasets respectively.

There was a clear trend between XY image resolution and segmented mask volume and length. This trend led to the under-segmentation of low-resolution TOF MRA images within the IXI and MIDAS repository datasets creating large neurovascular masks containing additional peripheral vessels irrelevant for aneurysmal identification such as venous sinuses and torcula. An exponential relationship between XY image resolution and the *k*_*T*_ thresholding parameter was defined to normalise segmentation across resolutions (Figure 5, Figure S2). Without normalisation, images from MIDAS and IXI repositories were significantly different to RBWH with respect to mask length and volume. After normalisation, mask lengths were more equal between repositories.

**Figure 5:**
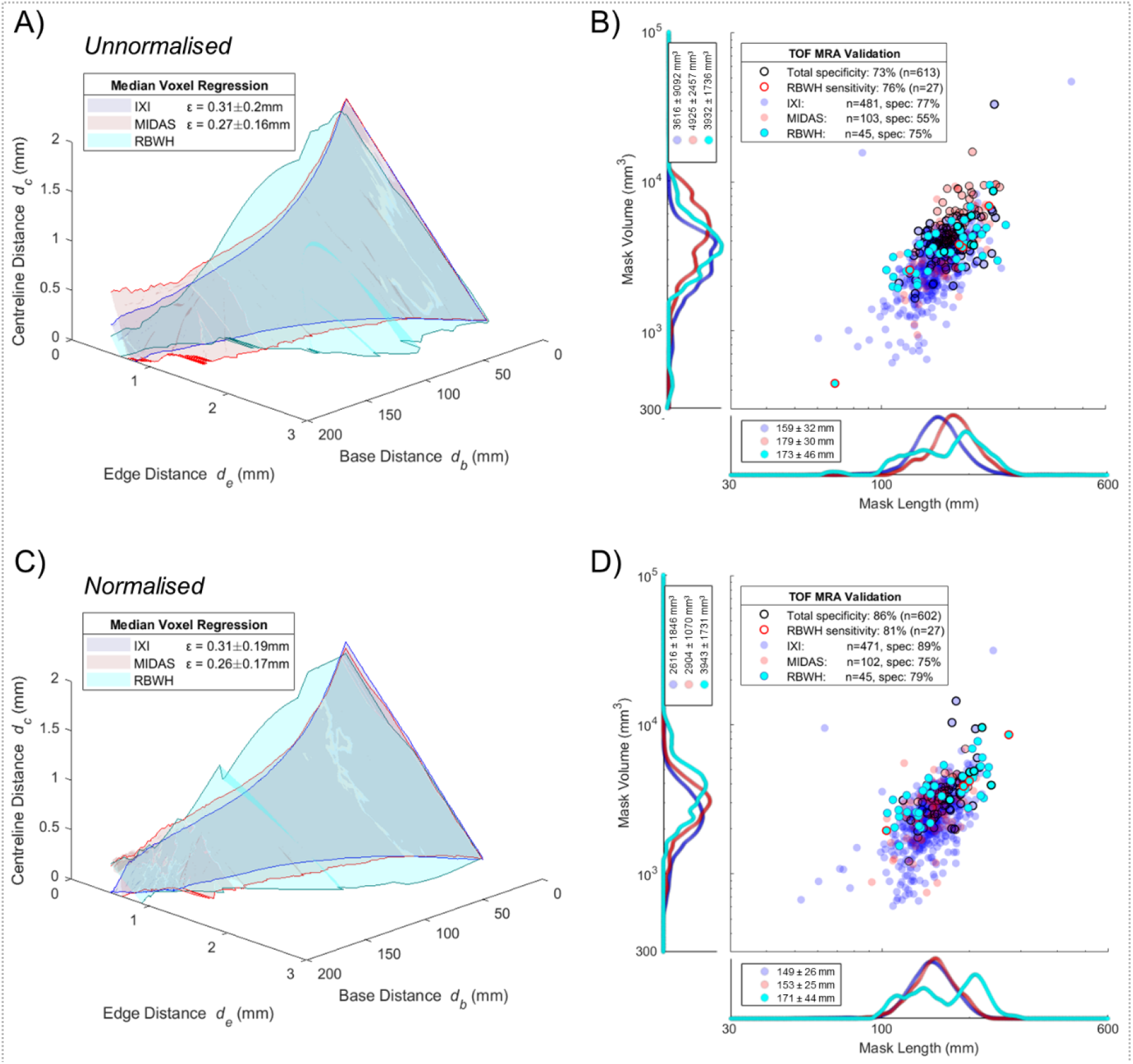
Algorithm validation and normalisation. (A,C) Median *h*(*d*_*e*_,*d*_*b*_) regressions of single-voxel morphometry for IXI (N = 571), MIDAS (N = 107) and RBWH (N = 45) repositories, where spatially-clustered voxels beyond a 96% confidence interval from individual regressions were considered UIA candidates. (B,D) Specificity and sensitivity were evaluated with false-positives circled black and false-negatives circled red, where errors existed predominately in large and long neurovascular masks. A trend between low resolution MRAs producing large segmented masks was normalised from (A,B) *k*_*T*_ = 1.78 by applying (C,D) 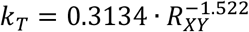, as detailed in Figure S2.

The algorithm’s identification achieved 81% sensitivity and 86% specificity, correctly identifying UIAs within 17 of 21 TOF MRAs within the RBWH database, and correctly identifying no aneurysms within 518 of 602 normal patient images within RBWH, IXI, and MIDAS databases (Figure 5). Nearly 80% of the 84 false positive UIA locations existed within the carotid siphon, where several tortuous bends deviate from normal cylindrical vessel geometry and poor contrast or segmentation could appear as though the siphon self-intersects (Bogunović et al., 2012; Duan et al., 2019). Most other false positive UIAs appeared within small overlapping peripheral arteries, especially for low-resolution images, such as the M3 and M4 segments in the anterior cerebral artery (as illustrated in Figure S3).

### 3.2. Detection pipeline is versatile to imaging resolution and MRA, CTA, or DSA modalities

This algorithm was principally developed for TOF MRA imaging, which represents a promising technique to 3D image intracranial vasculature without the use of intravenous contrast or x-ray radiation. However, many patients are unable to be imaged via MRA, including those who have previously had aneurysmal interventions with metallic neurosurgical clips or endovascular coils. Higher-resolution CTA and DSA are frequently employed as a primary or secondary imaging method to confirm aneurysm location and shape. In our RBWH MRA datasets, 82% of aneurysm-harbouring patients also had CTA or DSA imaging performed.

This algorithm was applied to abnormal and normal CTA and DSA images in Figure 6. These 3D images frequently have much higher XY resolution (0.15 - 0.30 mm/voxel) but lower Z resolution (1.0 - 2.0 mm/voxel) which can cause artefacts for vessels coarsely resolved in the Z-dimension. Even so, the algorithm was able to segment and identify UIAs within MRA, CTA, and DSA datasets, indicating its versatility across imaging modalities and clinical needs.

**Figure 6:**
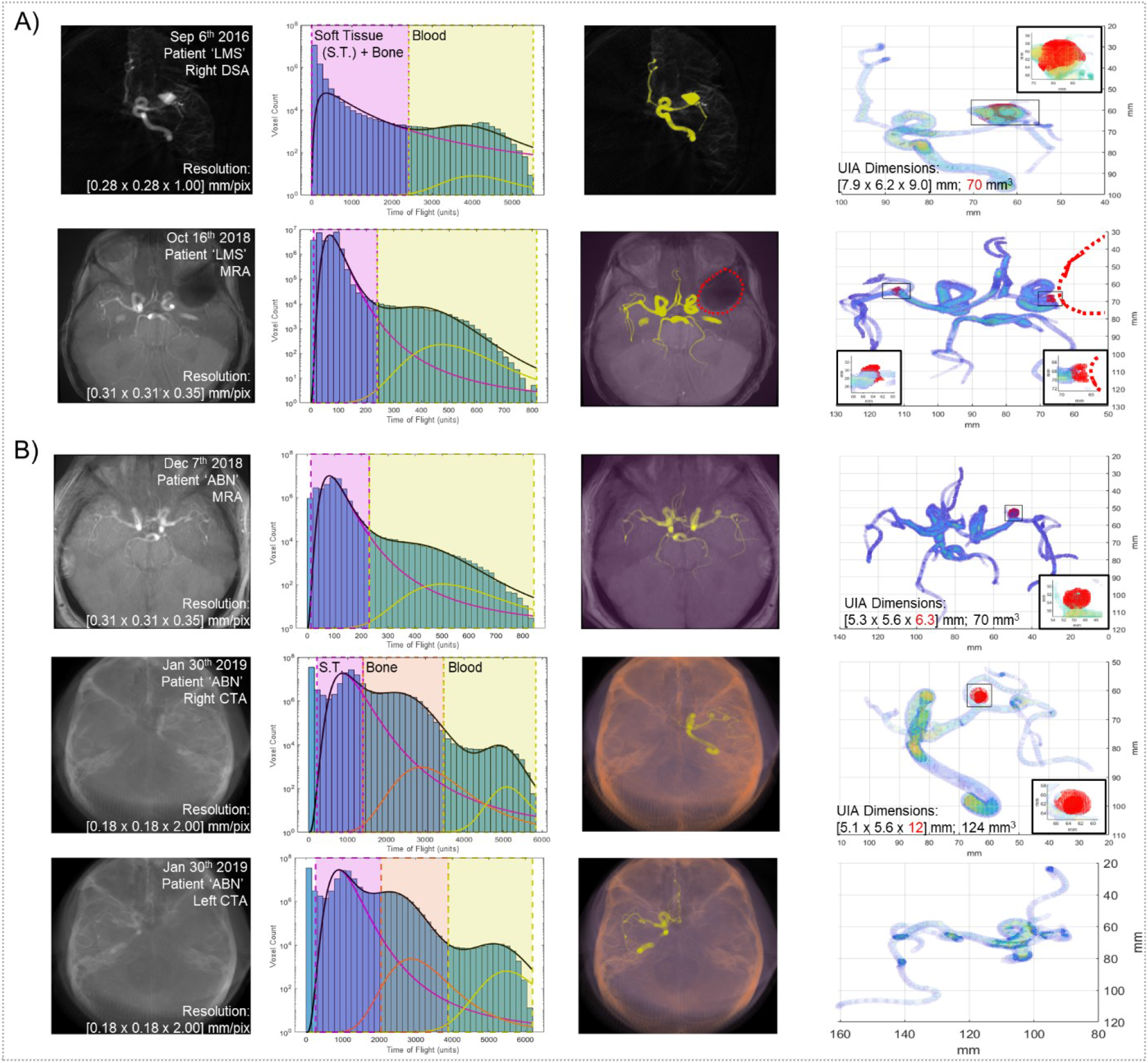
Algorithm versatility for different medical imaging modalities and resolutions. (A) Neurovasculature was imaged using right-hemisphere DSA (first row) before being rushed to surgery for endovascular coil placement. Later, this patient was imaged using TOF MRA with a large artefact at the location of the prior aneurysm (second row, red dotted circle). (B) Neurovasculature was imaged using TOF MRA or CTA for both brain hemispheres on the same date. Columns, from left to right, include original medical image mean intensity projection, voxel intensity histogram fit to the sum of 2 or 3 lognormal regressions, segmented soft tissue, bone, and/or vasculature masks, and mask voxel intensity heatmap with identified aneurysms. Measured UIA dimensions from CTA or DSA appear inaccurate, highlighted in red text. Abbreviation S.T. corresponds to ‘soft tissue’.

### 3.3. Adjustable detection sensitivity can identify early budding aneurysms

The algorithm can be tailored to suit specific clinical needs by adjusting two sensitivity parameters: the error threshold (*E*_*out*_) and the size threshold (*V*_*min*_) which identify outliers as candidate aneurysm regions. During algorithm validation and in Figure 5, these sensitivity parameters were kept constant across the 3 patient datasets and more than 700 patient images. However, sensitivity can be improved at the cost of specificity (false-positive aneurysm detection) by decreasing the minimum error or cluster size of candidate aneurysm regions to highlight small or slightly bulging intracranial vessels. While these bulging regions may be false positives generated by atypically tortuous neurovasculature or imaging or segmentation artefacts, lowering these detection thresholds could be useful to suggest potential small, difficult-to-spot, or secondary UIAs for radiological assessment.

To demonstrate the utility of this algorithm’s tuneable sensitivity, we identified a patient harbouring an UIA which was monitored over 5 TOF MRA imaging sessions between 2012 and 2018. During the initial visit in 2012, a large 4 × 3 mm saccular aneurysm was discovered on the left peripheral communicating artery and was monitored over the following 6 years. In 2018, a second bilateral aneurysm was discovered which led to a surgical decision of intervention. Using our algorithm at Figure 5’s validation sensitivity, we identified the same UIAs at the same timepoints as the clinicians identified. We then repeated our algorithm using a heightened sensitivity which detected abnormal voxel clusters larger than 0.7 mm^3^. Using this sensitivity, we could observe the growth of a small bulging region (0.9 – 2.9 mm^3^) at the same location that the bilateral aneurysm would form 6 years earlier, with no other false-positives detected (Figure 7). While such small regional voxel abnormalities may often occur due to imaging artefact or normal variances within neurovascular anatomy, such a sensitive identification could identify regions of radiologic interest similar to those performed in recent computer-aided diagnosis approaches (Faron et al., 2019; Miki et al., 2016; Nakao et al., 2018; Stember et al., 2019; Ueda et al., 2019).

**Figure 7:**
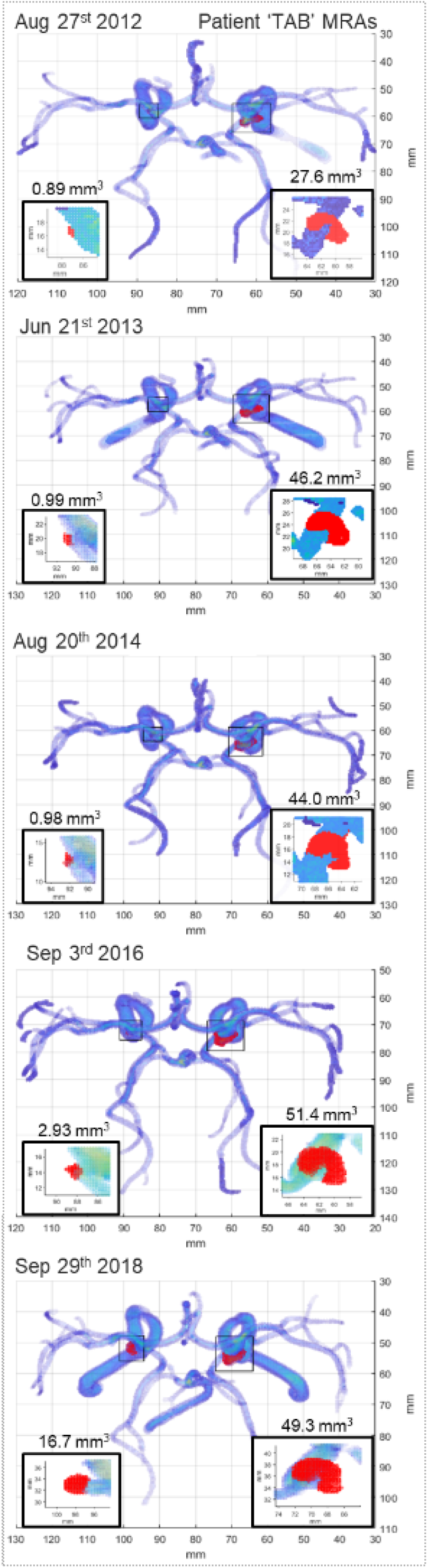
Single-case longitudinal study of an emerging second UIA. Patient neurovasculature was imaged using TOF MRA over five monitoring sessions across six years. The initial four sessions monitored any shape changes from a large burst saccular aneurysm within the right hemisphere (50 mm^3^ final size). On the final monitoring session, a second bilateral aneurysm was detected within the left hemisphere (20 mm^3^ final size) which led to the decision to operate. Implementing the ‘mask refinement’ option and an UIA detection sensitivity of 0.7 mm^3^, the emergence of a bulging region was detected in the location of the second aneurysm up to 6 years (4 imaging appointments) prior to its clinical identification by a radiologist.

Previous computer-aided diagnoses have reached high levels of sensitivity but have done so exhibiting very low specificity, incorrectly detecting several false-positive UIAs per image. These approaches still require substantial radiological interpretation to exclude these false-positives and may not improve the number of medical images a radiologist can assess within a certain amount of time. Furthermore, detection rates vary between radiologists and neuroradiologists and our approach enables an unbiased detection and characterisation of UIAs at mm^3^ resolution (Okahara et al., 2002). This user-adjustable approach will better provide both high-specificity screening of UIA presence and high-sensitivity UIA characterisation toward reduced radiologist workloads.

## 4. Discussion

We propose a new method to identify UIAs within 3D medical angiograms, principally within widely-used TOF MRAs (Thompson et al., 2015). The key innovations of our method include high sensitivity with high specificity and user-defined versatility while utilising large 3D medical images independent of interpatient comparison. Our method reaches at least an 81% sensitivity and 86% specificity, on-par with conventional computer aided MRA diagnoses achieving up to 83.6% sensitivity and 75% specificity, while analysing MRAs 10-fold faster (Miki et al., 2016; Štepán-Buksakowska et al., 2014; Yang et al., 2011). Sensitivities above 90% have been reached with other machine learning approaches but only while generating as many as 4 - 22 false-positives per image indicating specificities near 0% (Faron et al., 2019; Nakao et al., 2018; Shi et al., 2020; Stember et al., 2019; Ueda et al., 2019). One recent approach identified UIAs inside small ~30mm vessel segments with 88.5% sensitivity and 98.5% specificity, but required 8 hours to manually segment each neurovascular model into pieces (Yang et al., 2020). These machine learning approaches rely on large training datasets which may be difficult to acquire and time-consuming to train for 3D angiograms of incidentally-discovered UIAs. Altogether, computer-aided diagnosis have only increased radiologist diagnosis sensitivity from 64% to 69% and have not reduced radiologist interpretation time (Miki et al., 2016; Z. Shi et al., 2020; Štepán-Buksakowska et al., 2014).

Our clustered anomaly detection method presented in this paper calculates a unique morphometry regression for each individual image, and is not reliant on a large dataset of training images typical of current machine learning approaches (Faron et al., 2019; Miki et al., 2016; Štepán-Buksakowska et al., 2014; Yang et al., 2011). And while current approaches frequently analyse medical images of only one single imaging modality and resolution, our method was evaluated over 10 different TOF MRA resolutions from 5 different hospitals and also applied to several CTA and 3D DSA images. A specific utility of our method is its ability to tailor mask segmentation and UIA detection sensitivity depending on user demand. If high sensitivity at the cost of low specificity is preferred, the UIA detection threshold can be lowered. This could also assist with the detection of secondary UIAs or abnormal budding regions as demonstrated in Figure 7. While our method could mimic current approaches by determining one average morphometry regression across our large healthy dataset of images, such a method would be limited due to the anatomical complexity and variability common to intracranial angiograms. This variability can be reduced through image normalisation approaches, but a recent neurovascular atlas (544 healthy TOF MRAs) indicates MRA normalisation may only allow consistent segmentation for major arteries (Mouches and Forkert, 2019). Our approach has several limitations. It relies on the construction of an interconnected centreline throughout all vessels, which occasionally cannot be achieved due to TOF MRA bleb artefacts (Corfield et al., 2010; Mair, 2015). Furthermore, TOF MRA imaging achieves poorer resolution than CTA or DSA methodologies (Lin et al., 2018), limiting the detection of small aneurysms in peripheral neurovasculature. Detection sensitivity increased with local image resegmentation and would likely further increase with enhanced centreline estimation, image normalisation, or local polynomial regression or smoothing algorithms (Kerrien et al., 2017; Pelka et al., 2017; Wong and Chung, 2007). We prototyped several such developments which can improve sensitivity or specificity slightly but require substantial increases in processing time. Finally, our large TOF MRA dataset is heavily biased toward healthy patients. It will be necessary to recruit additional patient cases harbouring rare incidentally discovered UIAs in order to have greater confidence in our detection sensitivity.

The detection of UIAs prior to rupture allows for careful management to avoid haemorrhage. Fortunately, the incidental discovery of UIAs is becoming more frequent due to the increased use and resolution of MRA, a neurovascular imaging technique which does not require intravenous contrast or x-ray radiation (Thompson et al., 2015). While MRA may be a promising angiography technique to screen for UIAs in patients with a strong family history or those presenting migraines (Micieli and Kingston, 2019), it would be laborious, expensive, and unfeasible to engage expert neuroradiologists to review large numbers of cranial angiograms within a publicly-funded clinical imaging department (Z. Shi et al., 2020). In addition, once a UIA is discovered the surgical decision-making process remains ‘complex and controversial’ where as many as 58.3% of UIA patients undergo neuro or endovascular surgery (International Study of Unruptured Intracranial Aneurysms (ISUIA), 2003). While rupture risk is associated with UIA size and location aspect ratio, neck-to-body ratio, and intra-UIA fluid dynamics and wall thickness (Duan et al., 2018; Ishibashi et al., 2009; Russell et al., 2013), no widely-accepted prediction of rupture exists to guide surgical decision-making, and recent studies suggest decision-making has favoured interventional methods. Automated and rapid computational analyses of cranial angiograms could enable an unbiased screening of patient neurovasculature for UIAs and future identification of morphometric or blood flow features correlating to future UIA size and shape changes, surgical decisions, or rupture risk, or assessment of other vascular malformations (Chien et al., 2020; Huang et al., 2013; Z. Shi et al., 2020).

## 5. Conclusion

We have developed a rapid and automated 3D cranial angiogram analysis algorithm which segments neurovasculature, assesses single-voxel relationships between arterial morphometries, and identifies spatially-clustered voxel outliers as a potential UIA candidate. This method represents a significant improvement due to its high specificity and sensitivity, its independence from inter-image comparisons, and its versatility to imaging resolution and modality. While time-intensive conventional image analyses and training-intensive machine learning approaches can only achieve sensitivity above 80% with low specificity, our rapid automated method achieves 86% specificity with 81% sensitivity which reduces radiologist burden in assessing algorithm false-positives. This computational tool can serve as a second set of eyes to aid radiologist interpretation during UIA screening and has future value in morphometry-based rupture risk prediction.

All algorithms herein described to process and display medical images for UIA detection are freely available at https://github.com/mcallenby/UIAdetection2020.

## Acknowledgements

Our algorithms use rapid anisotropic Euclidean distance transforms generated by Yuriy Mishchenko (2015) and 3D medical image import algorithms by Dirk Jan Kroon (2011). We thank John Clouston for fruitful discussions. This work is supported by a RBWH Foundation Grant to CW, MP, JC, MCA and IHBI Inter-Theme Collaboration Grant to MCA and DAC. MCA is supported by an Advance Queensland Fellowship.

## CRediT author statement

**Mark C Allenby:** Conceptualization, methodology, software, formal analysis, validation, writing – original draft, writing – review & editing, visualization, funding acquisition. **Ee Shern Liang:** Resources, data curation, validation, writing – review & editing. **James Harvey:** Resources, data curation, validation, writing – review & editing. **Maria A Woodruff:** Supervision, project administration, writing – review & editing. **Marita Prior:** Resources, data curation, supervision, project administration. **Craig Winter:** Resources, data curation, supervision, writing – review & editing, funding acquisition. **David Alonso-Caneiro:** supervision, writing – review & editing, funding acquisition.

## Supplemental Material

**Figure S1:**
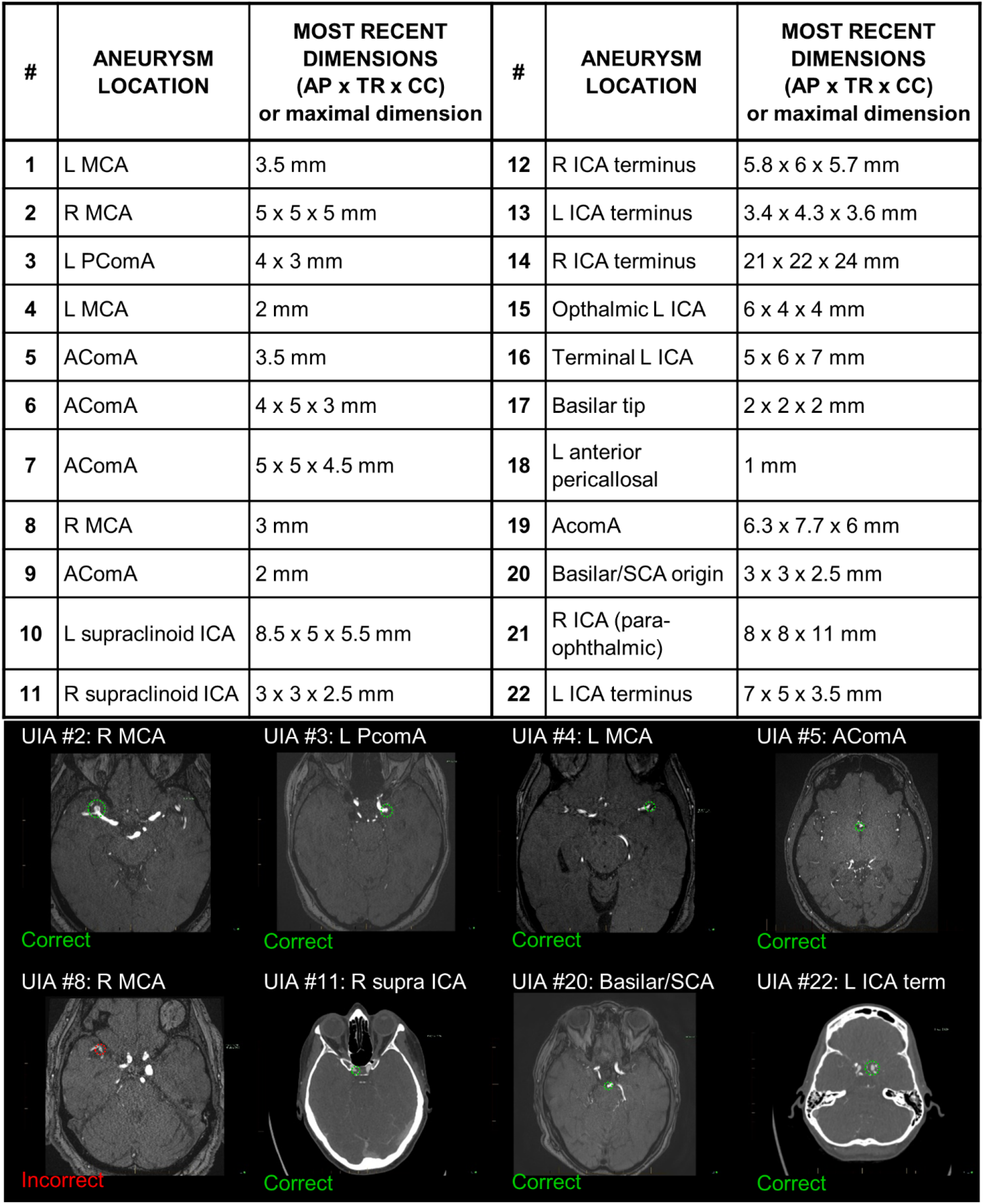
Radiologist-identified RBWH UIA sizes and locations. (top) De-identified table of representative UIA locations and measured dimensions where number corresponds to (bottom) radiologist annotated MRA or CTA images and the ability of our computational approach to correctly or incorrectly detect UIAs. Annotations include: L/R, left/right; MCA, middle carotid artery; PComA, peripheral communicating artery; AComA, anterior communicating artery; ICA, internal carotid artery; SCA, subclavian artery; AP, anteroposterior measurement; TR, traverse measurement; CC, craniocaudal (coronal) measurement.

**Figure S2:**
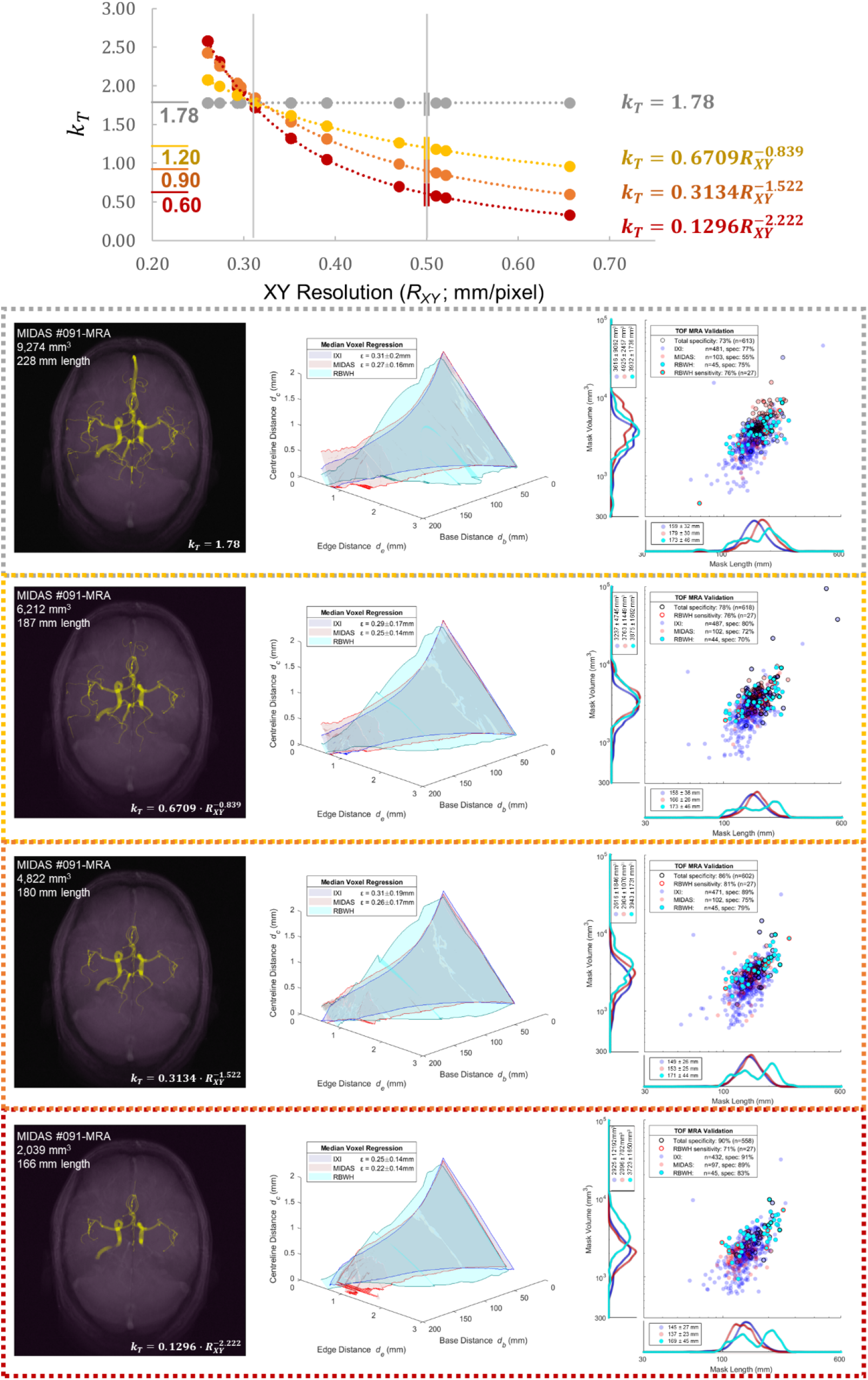
Impact of *k*_*T*_ normalisation on *R*_*XY*_ during validation and assessment. The constant *k*_*T*_ = 1.78 appeared ideal for images with XY resolution of *R*_*XY*_ = 0.32 mm/voxel but incorrectly segmented images with poorer *R*_*XY*_ near 0.5 mm/voxel. Power regressions are compared which normalise *k*_*T*_ values to 1.2 (yellow), 0.9 (orange), and 0.6 (red) at a *R*_*XY*_ of 0.5 mm/voxel while maintaining *k*_*T*_ = 1.78 at *R*_*XY*_ = 0.32 mm/voxel. Linear regressions at similar intercepts were also compared with poorer low *R*_*XY*_ image segmentations. Columns 2 and 3 of the grey and orange boxes are displayed in Figure 5.

**Figure S3:**
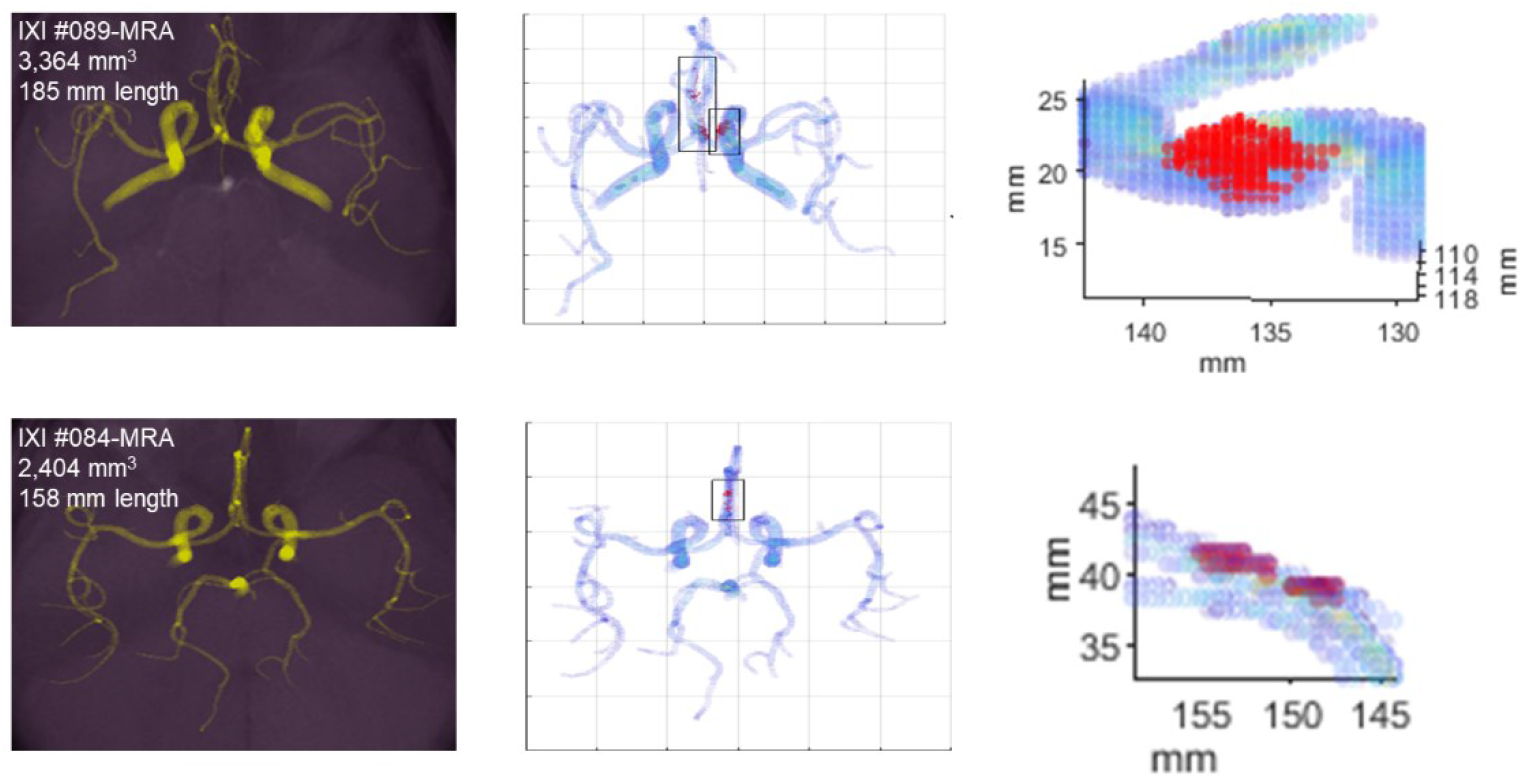
Examples of false-positive identifications in the carotid siphon (80% of false-positives; top row) and anterior cerebral artery (20% of false positives; bottom row) for poor resolution IXI repository TOF MRA images (0.47 × 0.47 × 0.8 mm/voxel). The carotid siphon anatomy rapidly changes diameter and angle in different ways for different patients, which produces false-positive detections of dissecting aneurysms (top right). The poor resolution images segment anterior cerebral arteries that comprise only a few voxels in diameter, and due to their low resolution, these arteries appear to overlap during segmentation, creating false-positive artefacts.

## References

Bogunović, H., Pozo, J.M., Cárdenes, R., Villa-Uriol, M.C., Blanc, R., Piotin, M., Frangi, A.F., 2012. Automated landmarking and geometric characterization of the carotid siphon. Med. Image Anal. 16, 889–903. https://doi.org/10.1016/j.media.2012.01.006

Chien, A., Callender, R.A., Yokota, H., Salamon, N., Colby, G.P., Wang, A.C., Szeder, V., Jahan, R., Tateshima, S., Villablanca, J., Duckwiler, G., Vinuela, F., Ye, Y., Hildebrandt, M.A.T., 2020. Unruptured intracranial aneurysm growth trajectory: Occurrence and rate of enlargement in 520 longitudinally followed cases. J. Neurosurg. https://doi.org/10.3171/2018.11.JNS181814

Corfield, L., Speirs, A., McCormack, D.J., Waltham, M., 2010. Time of Flight Magnetic Resonance Angiography: A Trap for the Unwary. EJVES Extra 19, e35–e37. https://doi.org/10.1016/j.ejvsextra.2010.01.002

Duan, H., Huang, Y., Liu, L., Dai, H., Chen, L., Zhou, L., 2019. Automatic detection on intracranial aneurysm from digital subtraction angiography with cascade convolutional neural networks. Biomed. Eng. Online 1–18. https://doi.org/10.1186/s12938-019-0726-2

Duan, Z., Li, Y., Guan, S., Ma, C., Han, Y., Ren, X., Wei, L., Li, W., Lou, J., Yang, Z., 2018. Morphological parameters and anatomical locations associated with rupture status of small intracranial aneurysms. Sci. Rep. 8, 1–7. https://doi.org/10.1038/s41598-018-24732-1

Faron, A., Sijben, R., Teichert, N., Freiherr, J., Wiesmann, M., Sichtermann, T., 2019. Deep learning-based detection of intracranial aneurysms in 3D TOF-MRA. Am. J. Neuroradiol. 40, 25–32. https://doi.org/10.3174/ajnr.A5911

Forkert, N.D., Illies, T., Moller, D., H., H., Saring, D., Fiehler, J., 2012. Analysis of the Influence of 4D MR Angiography Temporal Resolution on Time-to-Peak Estimation Error for Different Cerebral Vessel Structures. Am. J. Neuroradiol. 33, 2103–2109.

Huang, T.C., Chang, C.K., Liao, C.H., Ho, Y.J., 2013. Quantification of Blood Flow in Internal Cerebral Artery by Optical Flow Method on Digital Subtraction Angiography in Comparison with Time-Of-Flight Magnetic Resonance Angiography. PLoS One 8. https://doi.org/10.1371/journal.pone.0054678

International Study of Unruptured Intracranial Aneurysms (ISUIA), 2003. Unruptured intracranial aneurysms: natural history, clinical outcome, and risks of surgical and endovascular treatment. Lancet 362, 103–110.

Ishibashi, T., Murayama, Y., Urashima, M., Saguchi, T., Ebara, M., Arakawa, H., Irie, K., Takao, H., Abe, T., 2009. Unruptured intracranial aneurysms: Incidence of rupture and risk factors. Stroke 40, 313–316. https://doi.org/10.1161/STROKEAHA.108.521674

Jan Kroon, D., 2011. Matlab File Exchange “Read Medical Data 3D” [WWW Document]. URL https://au.mathworks.com/matlabcentral/fileexchange/29344-read-medical-data-3d

Jin, Z., Arimura, H., Kakeda, S., Yamashita, F., Sasaki, M., Korogi, Y., 2016. An ellipsoid convex enhancement filter for detection of asymptomatic intracranial aneurysm candidates in CAD frameworks. Med. Phys. 43, 951–960. https://doi.org/10.1118/1.4940349

Kerrien, E., Yureidini, A., Dequidt, J., Duriez, C., Anxionnat, R., Cotin, S., 2017. Blood vessel modeling for interactive simulation of interventional neuroradiology procedures. Med. Image Anal. 35, 685–698. https://doi.org/10.1016/j.media.2016.10.003

Leng, X., Wang, Y., Xu, J., Jiang, Y., Zhang, X., Xiang, J., 2018. Numerical simulation of patient‑specific endovascular stenting and coiling for intracranial aneurysm surgical planning. J. Transl. Med. 16, 1–10.

Li, M.H., Cheng, Y.S., Li, Y.D., Fang, C., Chen, S.W., Wang, W., Hu, D.J., Xu, H.W., 2009. Large-cohort comparison between three-dimensional time-of-flight magnetic resonance and rotational digital subtraction angiographies in intracranial aneurysm detection. Stroke. 40, 3127–3129. https://doi.org/10.1161/STROKEAHA.109.553800

Lin, A., Rawal, S., Agid, R., Mandell, D.M., 2018. Cerebrovascular Imaging: Which Test is Best? Clin. Neurosurg. 83, 5–18. https://doi.org/10.1093/neuros/nyx325

Mair, G., 2015. Lack of flow on time-of-flight MR angiography does not always indicate occlusion. BJR|case reports 2, 20150187. https://doi.org/10.1259/bjrcr.20150187

Mayo Foundation for Medical Education and Research, 2017. Patient Care & Helath Information: Brain Aneurysm & Carotid Artery Disease [WWW Document]. URL https://www.mayoclinic.org/diseases-conditions/brain-aneurysm/diagnosis-treatment/drc-20361595

Micieli, A., Kingston, W., 2019. An approach to identifying headache patients that require neuroimaging. Front. Public Heal. https://doi.org/10.3389/fpubh.2019.00052

Miki, S., Hayashi, N., Masutani, Y., Nomura, Y., Yoshikawa, T., Hanaoka, S., Nemoto, M., Ohtomo, K., 2016. Computer-assisted detection of cerebral aneurysms in MR angiography in a routine image-reading environment: Effects on diagnosis by radiologists. Am. J. Neuroradiol. 37, 1038–1043. https://doi.org/10.3174/ajnr.A4671

Mishchenko, Y., 2015. A fast algorithm for computation of discrete Euclidean distance transform in three or more dimensions on vector processing architectures. Signal, Image Video Process. 9, 19–27. https://doi.org/10.1007/s11760-012-0419-9

Mouches, P., Forkert, N.D., 2019. A statistical atlas of cerebral arteries generated using multi-center MRA datasets from healthy subjects. Sci. Data 6, 29. https://doi.org/10.1038/s41597-019-0034-5

Nakao, T., Hanaoka, S., Nomura, Y., Sato, I., Nemoto, M., Miki, S., Maeda, E., Yoshikawa, T., Hayashi, N., Abe, O., 2018. Deep neural network-based computer-assisted detection of cerebral aneurysms in MR angiography. J. Magn. Reson. Imaging 47, 948–953. https://doi.org/10.1002/jmri.25842

Nyúl, L.G., Udupa, J.K., Zhang, X., 2000. New variants of a method of MRI scale standardization. IEEE Trans. Med. Imaging. https://doi.org/10.1109/42.836373

Okahara, M., Kiyosue, H., Yamashita, M., Nagatomi, H., Hata, H., Saginoya, T., Sagara, Y., Mori, H., 2002. Diagnostic accuracy of magnetic resonance angiography for cerebral aneurysms in correlation with 3D-digital subtraction angiographic images: A study of 133 aneurysms. Stroke 33, 1803–1808. https://doi.org/10.1161/01.STR.0000019510.32145.A9

Pelka, O., Koitka, S., Johannes, R., Nensa, F., Friedrich, C.M., 2017. Intravascular Imaging and Computer Assisted Stenting, and Large-Scale Annotation of Biomedical Data and Expert Label Synthesis, MICCAI Workshop on Large-scale Annotation of Biomedical Data and Expert Label Synthesis (LABELS). https://doi.org/10.1007/978-3-319-67534-3

Russell, J.H., Kelson, N., Barry, M., Pearcy, M., Fletcher, D.F., Winter, C.D., 2013. Computational fluid dynamic analysis of intracranial aneurysmal bleb formation. Neurosurgery 73, 1061–1068.

Shi, Z., Hu, B., Schoepf, U.J., Savage, R.H., Dargis, D.M., Pan, C.W., Li, X.L., Ni, Q.Q., Lu, G.M., Zhang, L.J., 2020. Artificial Intelligence in the Management of Intracranial Aneurysms: Current Status and Future Perspectives. Am. J. Neuroradiol. 41, 373–379. https://doi.org/10.3174/ajnr.A6468

Shi, Z., Miao, C.C., Pan, C.W., Chai, X., Li, X.L., Xia, S., Gu, Y., Zhang, Y.G., Hu, B., Xu, W. Da, Zhou, C.S., Luo, S., Wang, H., Mao, L., Liang, K.M., Yu, Y.Z., Lu, G.M., Zhang, L.J., 2020. Clinically Applicable Deep Learning for Intracranial Aneurysm Detection in Computed Tomography Angiography Images: A Comprehensive Multicohort Study. medRxiv 2020.03.21.20040063. https://doi.org/10.1101/2020.03.21.20040063

Stember, J.N., Chang, P., Stember, D.M., Liu, M., Grinband, J., Filippi, C.G., Meyers, P., Jambawalikar, S., 2019. Convolutional Neural Networks for the Detection and Measurement of Cerebral Aneurysms on Magnetic Resonance Angiography. J. Digit. Imaging 32, 808–815. https://doi.org/10.1007/s10278-018-0162-z

Štepán-Buksakowska, I.L., Accurso, J.M., Diehn, F.E., Huston, J., Kaufmann, T.J., Luetmer, P.H., Wood, C.P., Yang, X., Blezek, D.J., Carter, R., Hagen, C., Hořínek, D., Hejčl, A., Rǒcek, M., Erickson, B.J., 2014. Computer-aided diagnosis improves detection of small intracranial aneurysms on MRA in a clinical setting. Am. J. Neuroradiol. 35, 1897–1902. https://doi.org/10.3174/ajnr.A3996

Thompson, B.G., Brown, R.D., Amin-hanjani, S., Broderick, J.P., Cockroft, K.M., Connolly, E.S., Duckwiler, G.R., Harris, C.C., Howard, V.J., Johnston, S.C.C., Meyers, P.M., Molyneux, A., Ogilvy, C.S., 2015. AHA / ASA Guideline Guidelines for the Management of Patients With Unruptured Intracranial Aneurysms. Stroke 46, 2368–2400. https://doi.org/10.1161/STR.0000000000000070

Ueda, D., Yamamoto, A., Nishimori, M., Shimono, T., Doishita, S., Shimazaki, A., Katayama, Y., Fukumoto, S., Choppin, A., Shimahara, Y., Miki, Y., 2019. Deep learning for MR angiography: Automated detection of cerebral aneurysms. Radiology 290, 187–194. https://doi.org/10.1148/radiol.2018180901

van Gijn, J., Kerr, R.S., Rinkel, G.J., 2007. Subarachnoid haemorrhage. Lancet 369, 306–318. https://doi.org/10.1016/S0140-6736(07)60153-6

Williams, L.N., Brown, R.D., 2013. Management of Unruptured Intracranial Aneurysms, in: Neurology: Clinical Practice. pp. 99–108.

Wong, W.C.K., Chung, A.C.S., 2007. Probabilistic vessel axis tracing and its application to vessel segmentation with stream surfaces and minimum cost paths. Med. Image Anal. 11, 567–587. https://doi.org/10.1016/j.media.2007.05.003

Yang, X., Blezek, D.J., Cheng, L.T.E., Ryan, W.J., Kallmes, D.F., Erickson, B.J., 2011. Computer-aided detection of intracranial aneurysms in MR angiography. J. Digit. Imaging 24, 86–95. https://doi.org/10.1007/s10278-009-9254-0

Yang, X., Xia, D., Kin, T., Igarashi, T., 2020. IntrA: 3D Intracranial Aneurysm Dataset for Deep Learning, arXiv.

